# Overriding defective FPR chemotaxis signaling in diabetic neutrophil stimulates infection control in diabetic wound

**DOI:** 10.1101/2021.09.09.459638

**Authors:** Ruchi Roy, Janet Zayas, Stephen J. Wood, Mohamed F. Mohamed, Dulce M. Frausto, Ricardo Estupinian, Eileena F. Giurini, Timothy M. Kuzel, Andrew Zloza, Jochen Reiser, Sasha H. Shafikhani

## Abstract

Infection is a major co-morbidity that contributes to impaired healing in diabetic wounds. Although impairments in diabetic neutrophils have been blamed for this co-morbidity, what causes these impairments and whether they can be overcome, remain largely unclear. Diabetic neutrophils, extracted from diabetic individuals, exhibit chemotaxis impairment but this peculiar functional impairment has been largely ignored because it appears to contradict the clinical findings which blame excessive neutrophil influx (neutrophilia) as a major impediment to healing in chronic diabetic ulcers. Here, we report that exposure to glucose in diabetic range results in impaired chemotaxis signaling through the FPR1 chemokine receptor in neutrophils, culminating in reduced chemotaxis and delayed neutrophil trafficking in wound in diabetic animals, and rendering diabetic wound vulnerable to infection. We further show that at least some auxiliary chemokine receptors remain functional under diabetic conditions and their engagement by the pro-inflammatory cytokine CCL3, overrides the requirement for FPR1 signaling and substantially improves infection control by jumpstarting the neutrophil response toward infection, and stimulates healing in diabetic wound. We posit that CCL3 may have real therapeutic potential for the treatment of diabetic foot ulcers if it is applied topically after the surgical debridement process which is intended to reset chronic ulcers into acute fresh wounds.

## Introduction

Diabetic foot ulcers are the leading cause of lower extremity amputations in the United States and are responsible for more hospitalizations than any other complication of diabetes (*1-5*). Infection with pathogenic bacteria, such as *Pseudomonas aeruginosa*, is a major co-morbidity that contributes to impaired healing in diabetic ulcers (*6-10*). Phagocytic leukocytes, particularly neutrophils (PMNs), play a major role defending wounds from invading pathogens (*11*). Neutrophil is the first inflammatory leukocyte that infiltrates into the wound (*12*). In addition to its antimicrobial functions mediated by phagocytosis, bursts of reactive oxygen species (ROS), antimicrobial (AMP) production, and neutrophil extracellular trap (NET) (*13, 14*), it also expresses various cytokines and chemokines that set the stage for the subsequent inflammatory and non-inflammatory responses, which further contribute to infection control and partake in healing processes (*15-19*). There appears to be a disconnect in that diabetic ulcers suffer from persistent non-resolving inflammation - characterized by increased neutrophils - yet they fail to control infection. Bactericidal functional impairments in diabetic neutrophils (PMNs) is thought to underlie defective infection control in diabetic wound (*20, 21*). What causes these impairments in diabetic neutrophils remains poorly understood, although the impairment severity has been associated with the degree of hyperglycemia (*20*), suggesting that exposure to high glucose levels may be to blame.

In addition to impaired bactericidal functions, diabetic neutrophils - (purified from the blood of diabetic patients) - also exhibit impaired chemotactic response (*22*). This peculiar functional impairment in diabetic neutrophils has not received much attention primarily because it appears to contradict the clinical findings which finds and blames excessive neutrophil response as a major impediment to healing in chronic diabetic ulcers (*23, 24*). Driven by this disconnect and in lieu of the fact that very little is known about neutrophil trafficking into diabetic wounds particularly early after injury and in response to infection, we sought to assess the possible impact of diabetic neutrophil chemotaxis impairment on the dynamics of neutrophil response and impaired infection control in diabetic wounds.

## Results

### Neutrophil trafficking is delayed in diabetic wounds

We generated full-thickness excisional wounds in db/db type 2 diabetic mice and their normal littermates C57BL/6, as described (*10, 25*), and challenged these wounds with PA103 *P. aeruginosa* bacteria (10^3^ CFU), which we have shown to establish a robust and persistent infection and cause wound damage in diabetic mice (*10*). Consistent with our previous report (*10*), db/db wounds contained 2-4 log orders more bacteria than normal wounds, indicating that diabetic wounds are vulnerable to increased infection (Fig. S1). We next collected wound tissues on days 1, 3, 6, and 10 post-infection and assessed them for their neutrophil contents by immunohistochemistry (IHC) using the neutrophil marker Ly6G (*26, 27*). Surprisingly, diabetic wounds exhibited substantially reduced neutrophil influx in diabetic wounds early after injury at days 1 and 3 but significantly higher neutrophil contents in day 6 and day 10 older diabetic wounds, as compared to normal wounds (Fig. 1a-b). Corroborating these data, myeloperoxidase (MPO) - (a marker for primarily activated neutrophils (*28*)) - was also substantially reduced in diabetic wounds early after injury at days 1 and 3 but significantly higher in day 10 wounds (Fig. 1c). Assessment of neutrophil contents in normal and diabetic infected day 1 wounds by flow cytometry - where neutrophils were identified as CD45^+^Ly6C/G^hi^CD11b^hi^ (*29, 30*) - further corroborated the inadequate neutrophil trafficking into diabetic wounds early after injury (Fig. 1d and Fig. S2). These data indicated that neutrophil response – (which is needed to combat infection) – is delayed in diabetic wounds, rendering these wounds vulnerable to infection early after injury.

**Fig. 1.**
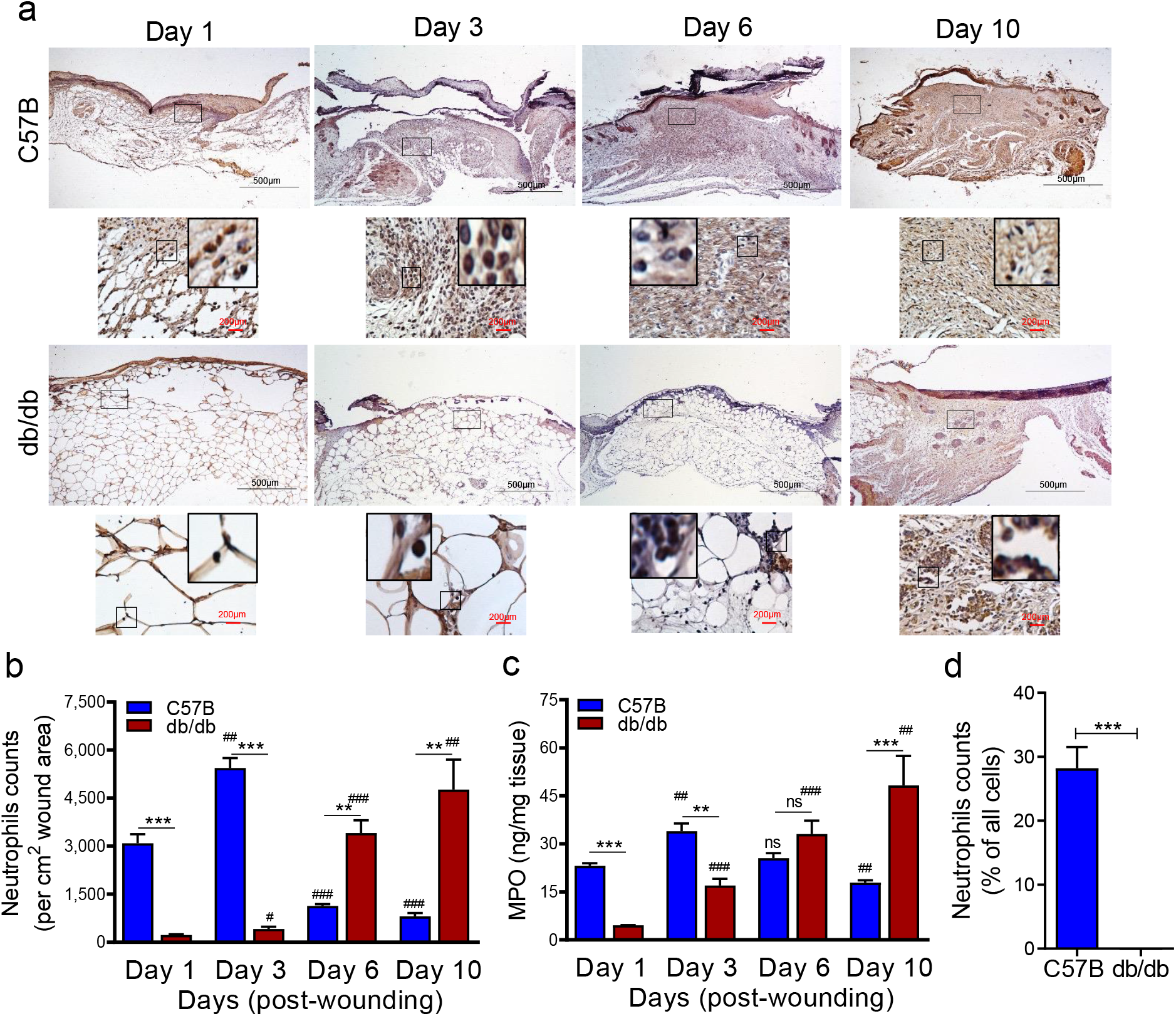
Neutrophil response is delayed in infected diabetic wound tissue. (a) Normal (C57B) and diabetic (db/db) wounds were infected with PA103 (10^3^ CFU). Wound tissues were harvested at indicated timepoints post-infection and assessed for neutrophil contents either by histological analysis using anti-Ly6G antibody (a-b), or by assessing MPO levels by ELISA (c). Representative regions from underneath the wounds extending in the dermis are shown at 40X and 400X magnification (top and bottom, respectively). A representative magnified region is also inserted in the 400X magnification images. Black scale bar = 500µm for 40x magnification and red scale bar = 200µm for 400x magnification. (d) Day 1 infected wound tissues of C57B and db/db were evaluated for their neutrophil response by flow cytometry. Corresponding data were plotted as the Mean ± SEM. (N=4; ns = not significant, *p<0.05; **p<0.01; ***p<0.001 – are comparisons made between C57B and db/db at indicated timepoints; or ^#^p<0.05; ^##^p<0.01; ^###^p<0.001 are comparisons made within each group to day 1 values respectively. Statistical analyses between groups were conducted by One-way ANOVA with additional post hoc testing, and pair-wise comparisons between groups were performed or by unpaired Student’s *t*-test).

### Chemotactic response through the FPR chemokine receptor is impaired in diabetic neutrophils

Depending on the tissue or the condition, neutrophil trafficking in response to injury and/or infection occurs in multiple waves mediated by ∼30 chemokine receptors on neutrophils and involves multiple signaling pathways (*31-37*). However, the initial neutrophil chemotaxis in response to injury or infection involves the activation of G protein–coupled formyl peptide chemokine receptors (e.g., FPR1) by *N*-formyl peptides, such as fMet-Leu-Phe (fMLF, a.k.a., fMLP), which is released either by injured tissues or by invading bacteria (*31, 38*). Activation of FPR receptors then leads to the upregulation and secretion of lipid signals, such as the leukotriene B_4_ (LTB_4_), which in turn activate BLT1, (another G protein-coupled receptor on neutrophils), amplifying neutrophil trafficking by enhancing the signaling through the FPR chemokine receptors (*36*). BLT1 activation in neutrophils by LTB_4_ also results in upregulation and secretion of pro-inflammatory cytokines, particularly IL-1β which in turn induces the expression and secretion of other chemokine ligands (i.e, CCL3 and CXCL1) in tissue resident epithelial cells and inflammatory leukocytes, which further amplify neutrophil trafficking and other inflammatory leukocytes including monocytes, by engaging their respective auxiliary chemokine receptors, such as CCR1 and CXCR2 (*36, 37, 39, 40*).

To assess the role of chemotaxis impairment in reduced neutrophil influx into diabetic wounds early after injury, we extracted neutrophils from blood of normal and diabetic mice and assessed chemotaxis signaling through FPR in response to fMLF. Compared to normal neutrophils extracted from C57B, db/db neutrophils were significantly impaired in their ability to chemotax toward fMLP (Fig. 2a). Consistent with reduced signaling through the FPR chemokine receptor, expression of FPR1 was significantly diminished in db/db neutrophils, as assessed by Western bloting (Fig. 2b-c). Further corroborating these data, the percentage of FPR1-positive neutrophils was significantly reduced in day 1 diabetic wounds, after accounting for reduced number of neutrophils in diabetic wounds early after injury by assessing equal number of neutrophils by flow cytometry (Fig. 2d).

**Fig. 2.**
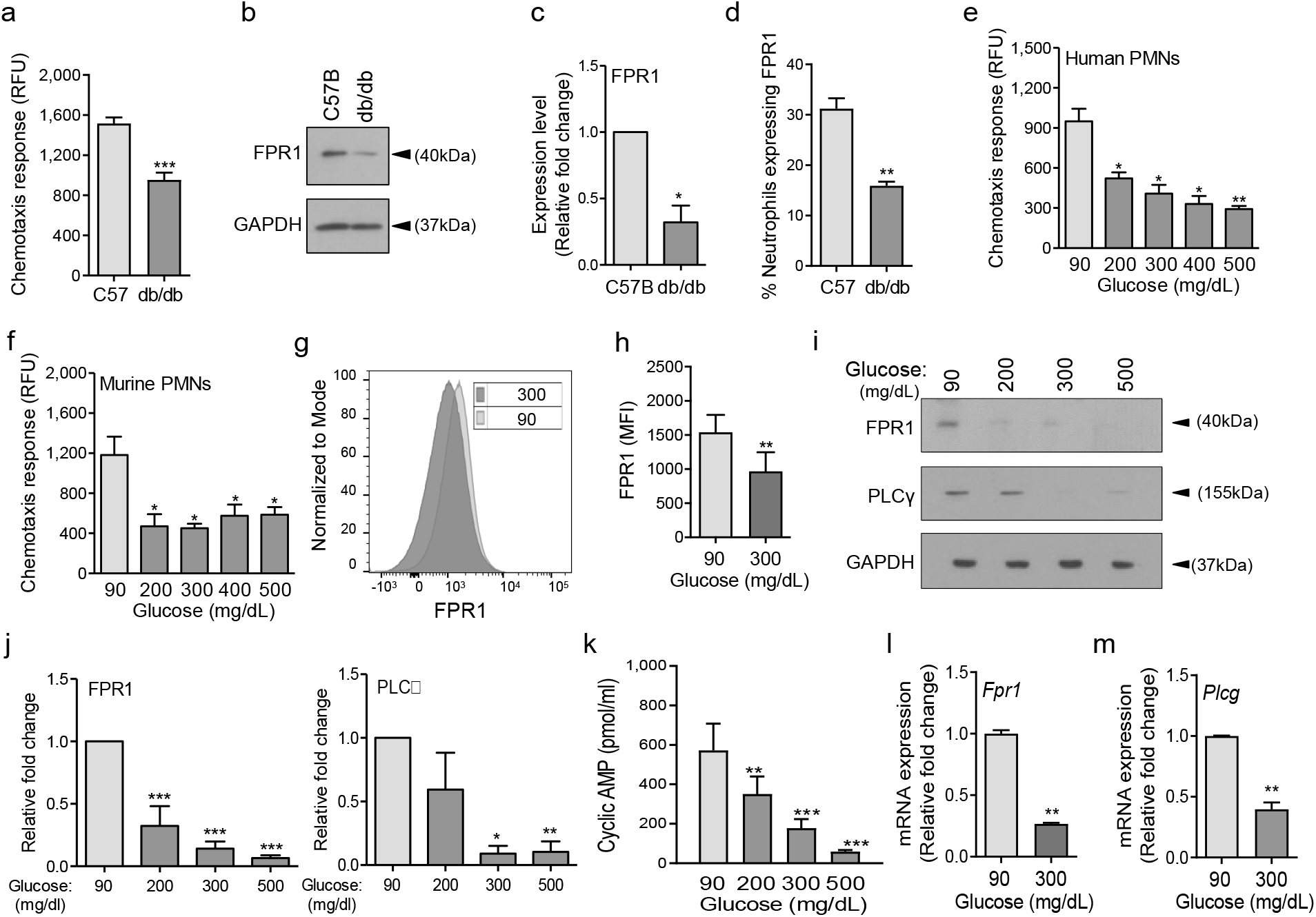
Chemotactic response is impaired in diabetic neutrophils through FPR1 chemokine receptor. (**a-b**) Neutrophils were extracted from the peripheral blood of C57B and db/db animals to assess their ability to chemotax toward 100nM fMLP (**a**), or for the expression of FPR1 by Western blotting (**b**). (**c**) Densitometry values associated with **(b)** are plotted as Mean ± SEM (N=4 mice/group). (**d**) Equal number of neutrophils (extracted from Day 1 C57B and db/db wounds) were assessed by for the expression of FPR1 on neutrophils by flow cytometry (N=4 mice/group). (**e-f**) Purified neutrophils from C57B bone marrow (**e**), or peripheral blood of non-diabetic individuals (**f**), were exposed to media containing glucose in normal (90 mg/dl) or diabetic range (200-500 mg/dl) for 1h to assess their ability to chemotax toward 100nM fMLP (N≥3 mice/group). (**g-m**) Neutrophils from C57B bone marrow were exposed to glucose in normal level (90 mg/dl) or in diabetic range (300 mg/dl) for 1h and assessed: for surface expression of FPR1 by flow cytometry **(g-h)**; for the expression of FPR1 and PLCγ by Western blotting (**i**) with corresponding densitometry values being plotted as Mean ± SEM (N=4 mice/group) in (**j-i**); or for mRNA expression analyses of FPR1 and PLCγ by RT-PCR in (**l-m**) (Data were plotted as the Mean ± SEM; N=2, each experiment repeated at least twice; ns = not significant, *p<0.05, **p<0.01, ***p<0.001; Student’s *t*-test); and for c-AMP levels in **(m)** (Data were plotted as the Mean ± SEM. (N≥3 mice/group, ns = not significant, *p<0.05, **p<0.01, ***p<0.001. Comparisons were made to the normal glucose level. Statistical analyses between groups were conducted by One-way ANOVA with additional post hoc testing, and pair-wise comparisons between groups were performed or by unpaired Student’s *t*-test).

Various studies have shown direct correlations between plasma glucose levels and prevalence and/or severity of infection in diabetic patients (*41-43*), suggesting that exposure to high glucose levels may be responsible for impaired neutrophil functions in diabetes. Consistent with these reports, short-term and long-term glycemic control in diabetic rats, significantly improved their ability to control *Staphylococcus aureus* infection (*44*). To assess the impact of high glucose on signaling through the FPR chemokine receptor, we purified neutrophils from human blood and C57B mice bone marrow (Fig. S4a-c and Materials & Methods), incubated them in media containing glucose in the normal range (90 mg/dl) or in the diabetic range (200-500 mg/dl) for 1h, and evaluated their chemotactic responses toward fMLF. (Of note, 1h exposure to high glucose in diabetic range had no effect on viability of neutrophils).

Exposure to high glucose levels caused significant reduction in chemotactic response to fMLF in both human and mouse neutrophils (Fig. 2e-f). While neutrophils exposed to normal glucose showed a bell-shaped curve in their chemotaxis response toward fMLF concentrations (0.01-1000nM) with 100nM being the optimum concentration, neutrophils exposed to high glucose showed flat chemotaxis response toward these fMLF concentrations, trending toward lower chemotaxis at higher concentrations (Fig. S4d), indicating that high fMLF ligand concentrations cannot rescue chemotaxis signaling through FPR1 receptor. The bell-shaped response to fMLF in normal neutrophils is in line with previous reports showing reduction in neutrophil chemotactic responses to other ligands at high concentrations (*45, 46*). Of note, exposure to high glucose also caused drastic reductions in FPR1 and PLCγ protein levels, as well as cAMP levels (Fig. 2i-k), which are all required to mediate FPR-mediated chemotaxis in neutrophils (*36, 47, 48*). Corroborating these data, exposure to high glucose resulted in significant reductions in the FPR1 and PLC_γ_ transcription as determined by mRNA analysis by RT-PCR (Fig. 2l-m). Collectively, these data indicated that elevated glucose levels in diabetes is responsible for the reduced chemotactic response through the FPR1 chemokine receptor in diabetic neutrophils.

### Some auxiliary chemokine receptors remain functional under diabetic conditions

Although, the initial neutrophil chemotactic response through the FPR receptors and the amplification of neutrophil chemotactic responses via other auxiliary chemokine receptors are interconnected and occur sequentially *in vivo* (*32-37*), none of these receptors appear to be essential on their own and their defects can be overcome by engaging other receptors (*37, 49, 50*). Chronic diabetic ulcers suffer from increased neutrophil contents (*23, 24*), indicating that diabetic neutrophils are capable of migrating into the wound, albeit at dysregulated kinetics as our data show (Fig. 1). Together, these findings suggested that chemotactic responses of diabetic neutrophils - though impaired through the FPR1 receptor (Fig. 2) - may be functional through one or more auxiliary chemokine receptors that mediate the amplification phase of neutrophil trafficking in wound and toward infection.

To evaluate this possibility, we assessed chemotactic responses toward CCL3 in human and mouse neutrophils after 1h exposure to glucose at normal or diabetic levels. The reason we focused on CCL3 was because it activates multiple auxiliary chemokine receptors, namely CCR1, CCR4, and CCR5 (*51-53*). Of note, CCR1 is an important chemokine receptor that is implicated in neutrophil trafficking to post-ischemic tissues (*54*) and ischemia is an important co-morbidity associated with impaired healing in diabetic wound (*2, 55*). Data indicated that exposure to glucose in the diabetic range did not affect the chemotactic responses toward CCL3 in human or mouse neutrophils (Fig. 3a-b), suggesting that these auxiliary receptors are unaffected by high glucose. To corroborate these data, we assessed the impact of high glucose exposure on CCR1 auxiliary chemokine receptor. In line with chemotaxis data, CCR1 expression remained unaffected in neutrophils after exposure to high glucose as assessed by Western blotting (Fig. 3c-d), by mRNA analysis (Fig. 3e), and by surface expression analysis (Fig. 3f). Further corroborating these data, CCR1 expression was similar in neutrophils extracted from the blood of db/db and C57B mice (Fig. 3h-i), and the percentage of CCR1-positive neutrophils in db/db day 1 wounds were similar to C57B day 1 wounds, after accounting for the reduced number of leukocytes in day 1 diabetic wounds by assessing equal number of neutrophils by flow cytometry (Fig. 3j). Of note, surface expression of auxiliary chemokine receptor CXCR2, (another important auxiliary chemokine receptor involved in the amplification of neutrophil response in wound and toward infection (*31, 56*)), on neutrophils and chemotaxis through the CXCR2 in response to CXCL1 (a.k.a. KC) - a known ligand for CXCR2 (*57*) - were also unaffected by high glucose exposure in neutrophils (Fig. S5). Collectively, these data suggested that at least CCR1 and CXCL2 auxiliary chemokine receptors may remain functional under diabetic conditions.

**Fig 3.**
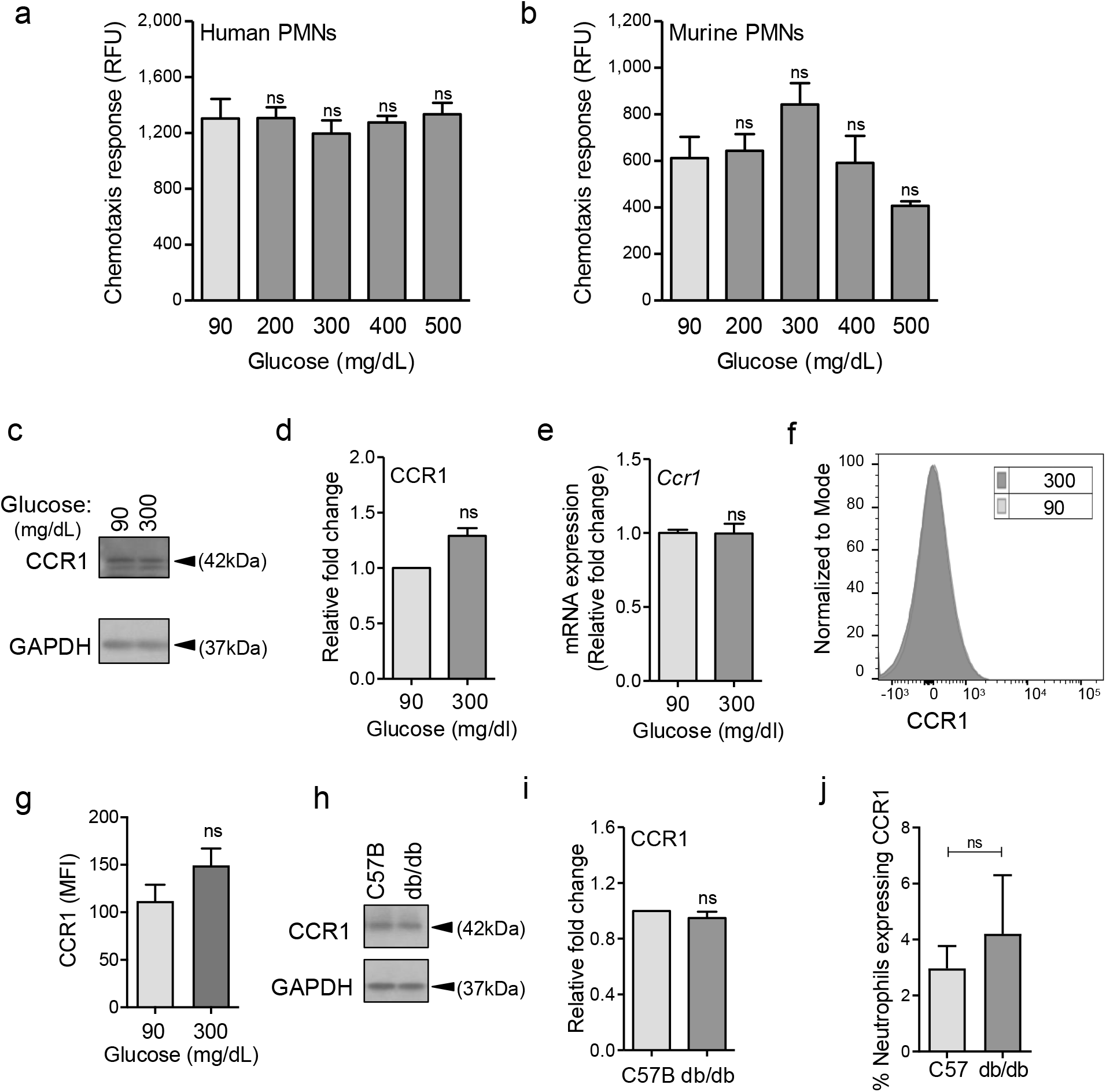
CCR1 receptor remains functional under diabetic conditions. Human **(a)** or mouse (**b**) neutrophils were examined for their chemotactic responses toward CCL3 (5ng/ml) after 1h exposure to glucose in normal (90 mg/dl) or diabetic range (200-500 mg/dl). (**c-g**) Neutrophils extracted from bone marrow of C57B were exposed to normal glucose (90 mg/dl) or high glucose (300 mg/dl) for 1h and assessed for CCR1 expression by Western blotting (c**-d);** for mRNA transcription analysis by RT-PCR (**e**); and for CCR1 surface expression **(f-g)**. (**h-i**) Neutrophils were purified from peripheral blood of normal (C57B) or diabetic (db/db) mice and assessed for the expression of CCR1 by Western blotting (**h**) and their respective densitometry values were plotted as the Mean ± SEM and shown in (**i**). (**j**) Equal number of neutrophils (extracted from Day 1 C57B and db/db wounds) were assessed by for the expression of CCR1 on neutrophils by flow cytometry (N=4; ns = not significant, *p<0.05, **p<0.01, ***p<0.001. Statistical analyses between groups were conducted by One-way ANOVA with additional post hoc testing, and pair-wise comparisons between groups were performed or by unpaired Student’s *t*-test).

### Topical treatment with CCL3 bypasses the requirement for FPR signaling and enhances neutrophil trafficking and infection control in diabetic wound

If CCL3 can rescue neutrophil chemotactic response in a situation where FPR signaling is impaired as our data indicate (Fig. 3a-b), why is neutrophil trafficking so severely diminished in diabetic wounds early after injury (Fig. 1). As discussed above, production of ligands for auxiliary chemokine receptors in tissue ultimately depends on FPR chemokine receptor activation (*36, 37, 39, 40*), suggesting that CCL3 expression may also be reduced in diabetic wounds early after injury. In line with this notion, CCL3 expression was substantially reduced in day 1 diabetic wounds, as assessed by mRNA analysis and Western blotting, after accounting for reduced number of leukocytes by normalizing the data with 18S or GAPDH loading control respectively (Fig. 4a-c). These data suggested that although auxiliary chemokine receptors on neutrophils may be functional in diabetic neutrophils, they may not be functioning in diabetic wounds early after injury because of inadequate ligands. If this is the case, augmenting diabetic wounds with CCL3 early after injury should be able to overcome deficiency in the FPR signaling and enhance neutrophil migration into diabetic wounds.

**Fig 4.**
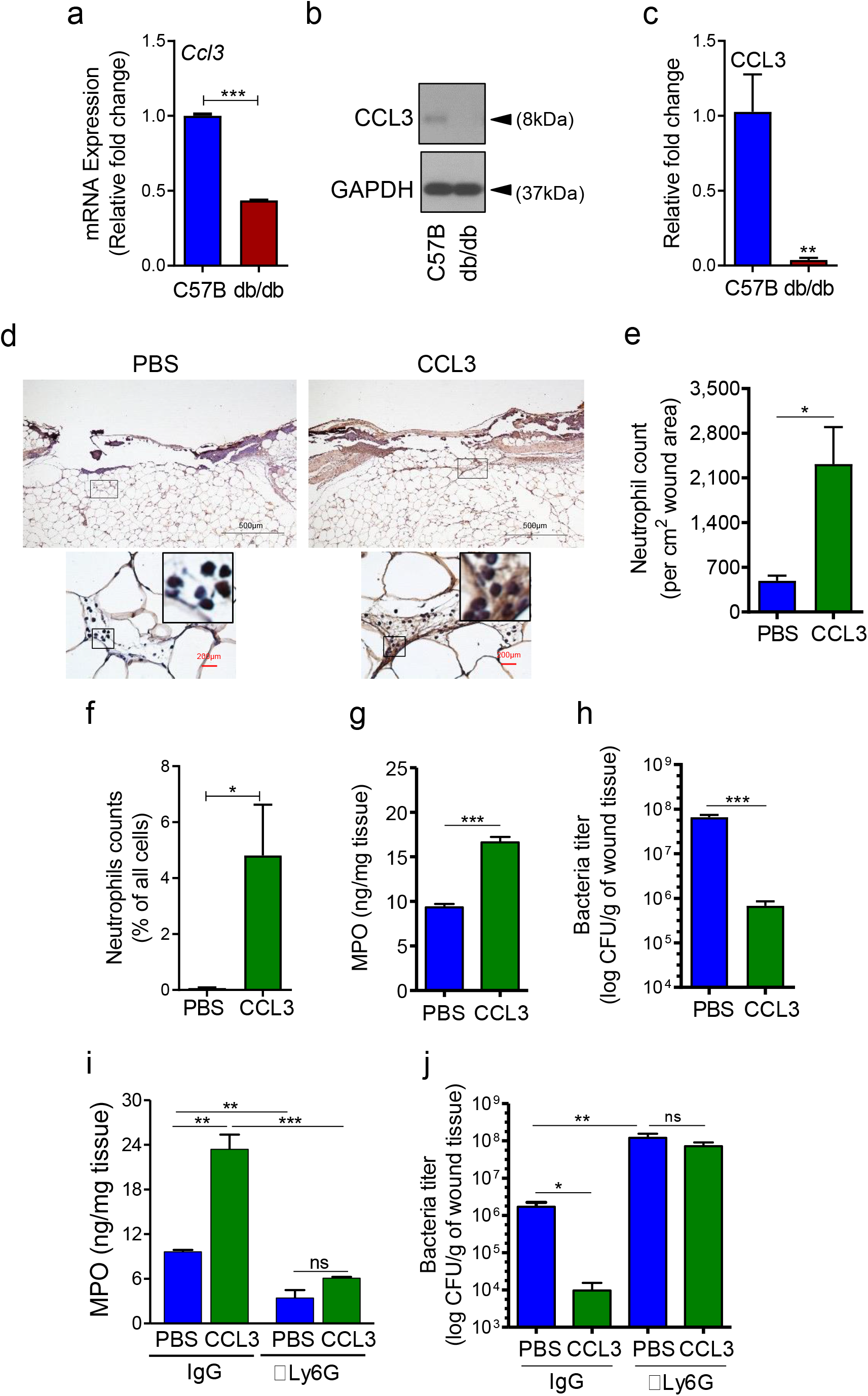
CCL3 topical treatment enhances neutrophil response and infection control in diabetic wound. **(a-c)** Day 1 wound tissues of C57B and db/db were harvested and assessed for the CCL3 mRNA levels by RT-PCR **(a)** and by Western blotting **(b-c)**, and the data were plotted as the Mean ± SEM (N≥4 mice/group), after normalization to 18S and GAPDH respectively, to account for reduced leukocytes in day 1 diabetic wounds. (**d-e)** db/db diabetic wounds were treated with either PBS or CCL3 (1μg/wound) and infected with PA103. 24h post-infection, wounds were collected and assessed for their neutrophil contents by histological analysis using a-Ly6G antibody. (**d**) Representative wound images at 40X and 400X magnification (top and bottom, respectively) are shown. Inserts are representative magnified regions within the 400X magnification images. Black scale bar = 500µm for 40x magnification and red scale bar = 200µm for 400X magnification. (**e**) Data are shown as Mean ± SEM (N≥4 mice/group, ≥9 random fields/wound/mouse). (**f**) Flow cytometry data showing increased neutrophil response in CCL3-treated Day 1 db/db infected wounds. (**g**) Neutrophil response was assessed for neutrophil marker MPO by ELISA. (**h-i**) db/db mice received either a-Ly6G (100μg/mouse) to cause neutrophil depletion or an IgG isoform as control, by intraperitoneal (i.p.) injection. 24h after injection, IgG or a-Ly6G-treated animals were wounded and treated with either PBS or CCL3 and infected with PA103. The impact of neutrophil depletion on the ability of CCL3 treatment to boost infection control in diabetic wound was assessed by MPO analysis (**i**) and CFU count determination (**j**) in day 1 wounds. Data were plotted as Mean ± SEM (N≥4 mice/group). (For all panels; ns = not significant, *p<0.05; **p<0.01, ***p<0.001. Statistical analyses between groups were conducted by One-way ANOVA with additional post hoc testing, and pair-wise comparisons between groups were performed or by unpaired Student’s *t*-test).

To test our hypothesis, we treated db/db wounds topically with CCL3 (1µg/wound) prior to infection and assessed its impact neutrophil response and infection control in diabetic wounds. One-time topical treatment with CCL3 significantly increased neutrophil trafficking in day 1 diabetic wounds, as assessed by Ly6G histological analysis (Fig. 4d-e), by flow cytometry (Fig. 4f), and by MPO analysis (Fig. 4g). Importantly, CCL3 treatment significantly enhanced the ability of diabetic wounds to control infection, as demonstrated by nearly a 2 log-order reduction in the number of bacteria contained in the CCL3-treated db/db wounds (Fig. 4h).

To assess the dependence enhanced infection control on neutrophils in CCL3-treated diabetic wounds, we depleted db/db mice of neutrophils by anti-Ly6G antibody (*58*), 24h prior to wounding and assessed the impact of neutrophil depletion on the ability of CCL3-treated db/db wounds to control *P. aeruginosa* infection. Anti-Ly6G reduced the neutrophil contents in circulation by ∼97% and in wound by ∼75% (Fig. 4i and Fig. S5a-b). Neutrophil-depletion resulted in 2 log-order more bacteria in diabetic wounds, indicating that despite their known bactericidal functional impairments (*20, 21*), diabetic neutrophils still contribute to infection control in these wounds (Fig. 4j). Importantly, neutrophil-depletion completely abrogated CCL3’s beneficial effects in boosting antimicrobial defenses against *P. aeruginosa* in diabetic wounds (Fig. 4j), indicating that CCL3-induced enhanced infection control in diabetic wound is completely dependent on its ability to enhance neutrophil response in diabetic wound.

### Treatment with CCL3 does not lead to persistent non-resolving inflammation in infected diabetic wounds and stimulates healing

Although, treatment with CCL3 substantially improved diabetic wound’s ability to control infection by enhancing neutrophil response toward infection early after injury in day 1 wounds (Fig. 4), it remained a possibility that CCL3 treatment could have long-term adverse consequences, as it could lead to heightened inflammatory environment which would be detrimental to the process of tissue repair and healing in diabetic wounds. Afterall, persistent non-resolving inflammation, (as manifested by increases in pro-inflammatory cytokines and neutrophils), is considered a major contributor to healing impairment in diabetic foot ulcers (*23, 24*).

We assessed the long-term impact of CCL3 treatment on neutrophil responses in diabetic wound. Data indicated that while neutrophils continued to rise in the mock-treated db/db infected wounds as they aged, in the CCL3-treated diabetic wounds, neutrophils were significantly higher during the acute phase of healing early after injury on days 1 and 3 but declined substantially in old wounds on days 6 and 10 (Fig. 5a-b).

**Fig. 5.**
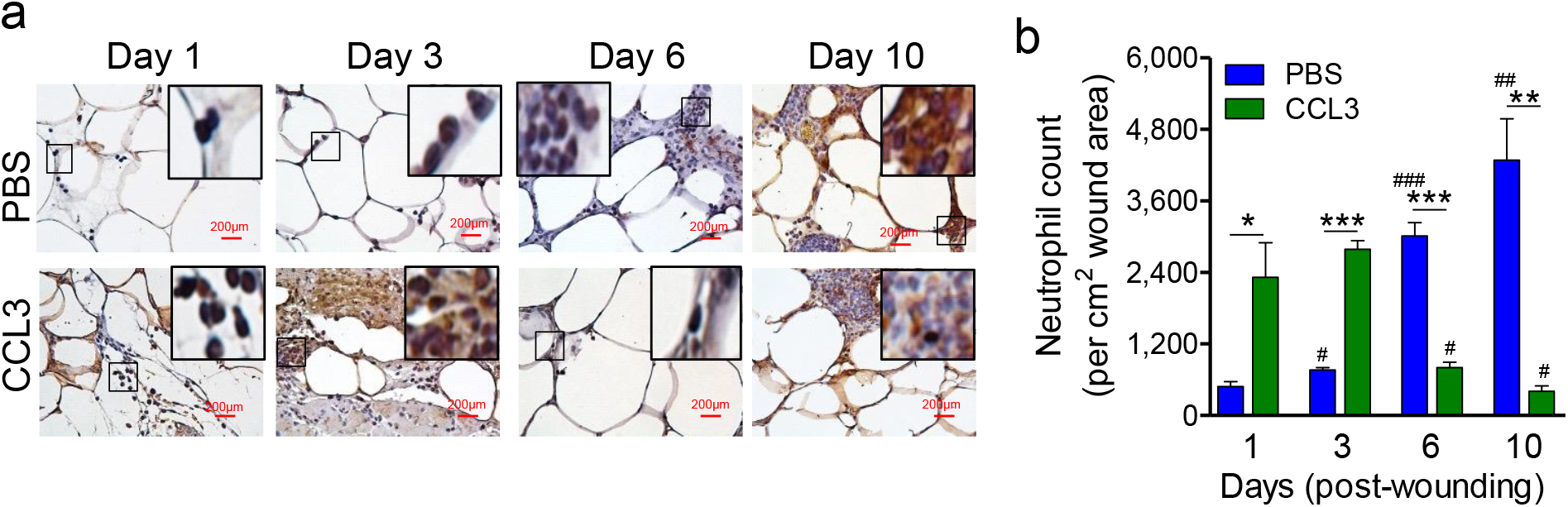
Treatment with CCL3 does not lead to persistent inflammation in infected diabetic wounds. db/db wounds were treated with PBS or CCL3 (1μg/wound) and infected with PA103 (1000 CFU). Wound tissues were collected at indicated timepoints and assessed for their neutrophils contents by histological analysis using neutrophil marker Ly6G staining. Representative images of regions from underneath the wounds extending in the dermis at 400X magnification are shown in (**a**). Representative magnified regions are inserted in the images. Red scale bars = 200μm. (Representative full wound images of these staining can be found in Fig. S7 and Fig. 6c). The corresponding data are plotted as the Mean ± SEM and are shown in (**b**). (N≥3 mice/group; ns = not significant; *p<0.05, **p<0.01, ***p<0.001, ^#^p<0.05, ^##^p<0.01, ^###^p<0.001. Statistical analyses between groups were conducted by One-way ANOVA with additional post hoc testing, and pair-wise comparisons between groups were performed or by unpaired Student’s *t*-test. (*) denotes significance between groups while (^#^) indicates significance within the same group in comparison to day 1 of respective wound groups).

Encouragingly, CCL3 treatment also significantly stimulated healing in infected diabetic wounds, as assessed by wound area measurement (Fig. 6a-b), while mock-treated diabetic wounds became exacerbated as the result of *P. aeruginosa* infection, as we had previously shown (*10*). Corroborating these results, CCL3-treated infected diabetic wounds were completely re-epithelized and exhibited epidermal thickening as assessed by H&E histological analysis, while mock-treated infected diabetic wounds became exacerbated (Fig. 6c-d).

**Fig 6.**
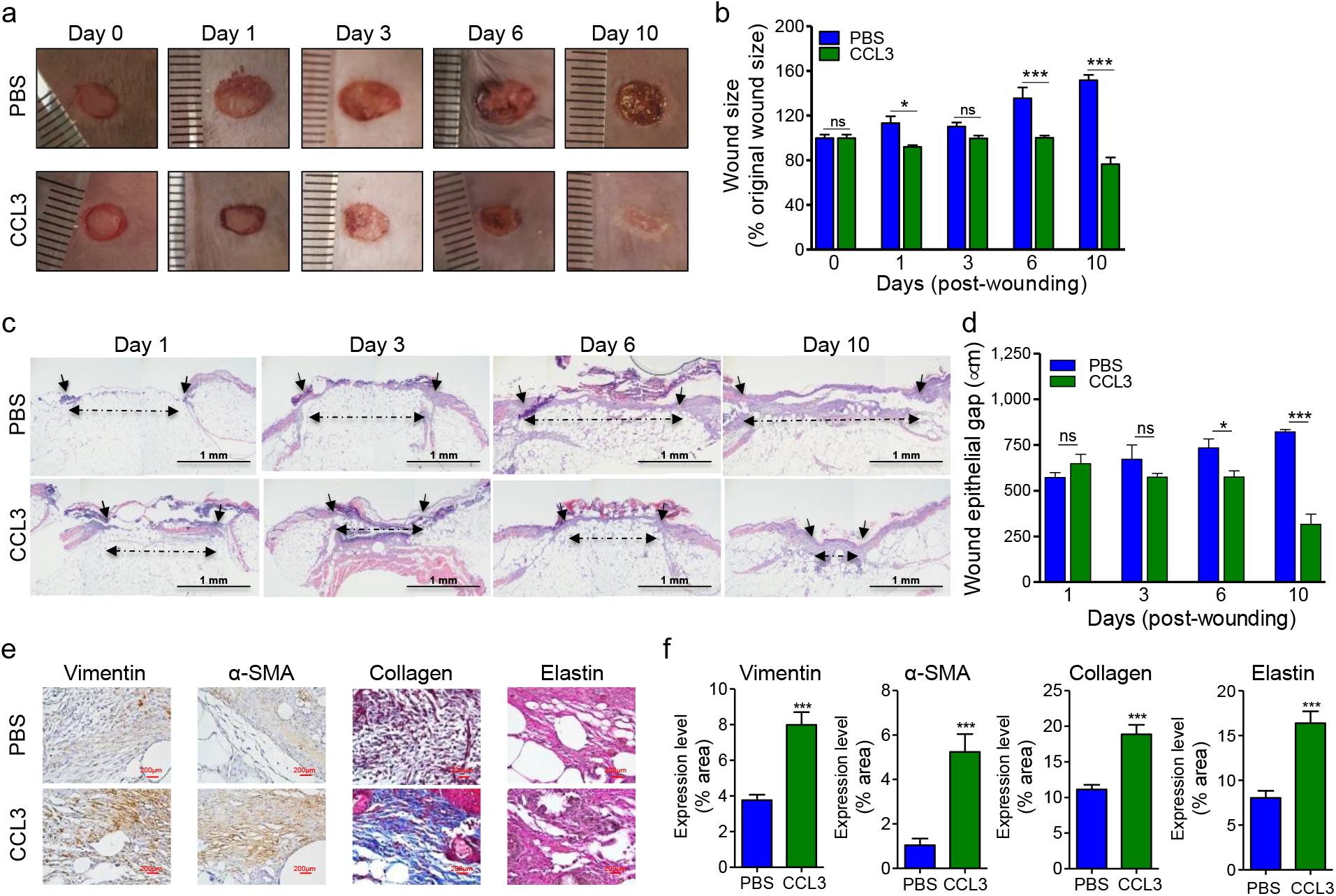
Treatment with CCL3 stimulates healing in infected diabetic wounds. **(a-d)** db/db wounds were either treated with PBS or CCL3 and infected with PA103 (1000 CFU). Wound healing was assessed at indicated timepoints by digital photography (**a**) or by H&E histological analysis of re-epithelization (**c**). Representative regions from underneath the wounds extending in the dermis are shown at 40X magnification are shown in (**c**). (Black scale bar = 1mm, and the wound gap is shown by dotted line). The corresponding data for (**a & c**) are shown in (**b & d**) as the Mean ± SEM. (N≥4 mice/group, ≥9 random fields/wound/mouse; ns = not significant; *p<0.05, **p<0.01, ***p<0.001, Student’s *t*-test). (**e-f**) Day 10 db/db wounds (treated with either PBS or CCL3 and infected with PA103) were assessed for fibroblast, myofibroblast, elastin and cartilage healing markers by vimentin, α-SMA, Masson’s Trichrome, and elastin staining respectively. **(e**) Representative regions from underneath the wounds extending in the dermis are shown at 400X magnification. (Red scale bar=200µm. For the corresponding full wound images at 40X magnification, see Fig. S8). (**f**) The corresponding data are plotted as the Mean ± SEM. (N=4 mice/group, ≥9 random fields/wound/mouse; ***p<0.001. Statistical analyses between groups were conducted by One-way ANOVA with additional post hoc testing, and pair-wise comparisons between groups were performed or by unpaired Student’s *t*-test).

Fibroblasts and myofibroblasts are key players in extracellular matrix production and granulation tissue maturation during the proliferation and the remodeling phases of wound healing (*59-61*). However, persistent inflammatory environment in diabetic wounds adversely impacts the functions of fibroblast and myofibroblast, culminating in reduced collagen and elastin extracellular matrix deposition and impaired healing in diabetic chronic wounds (*19, 62, 63*). *P. aeruginosa* infection further exacerbates inflammation and reduces collagen deposition in diabetic wounds (*10*). We evaluated the impact of CCL3 treatment on fibroblast, myofibroblast, collagen, and elastin in day 10 diabetic wounds, using their respective markers: Vimentin, α-SMA, Elastin, and Masson’s Trichrome staining (*10, 59, 64*). CCL3-treated wounds showed significant increases in all these healing markers (Fig. 6e-f and Fig. S7). Collectively, these data indicate that diabetic wounds are not destined to develop persistent non-resolving inflammation, provided that the dynamics of neutrophil trafficking is restored in these wounds early after injury.

## Discussion

Diabetic wounds are highly susceptible to infection with pathogenic bacteria, such as *P. aeruginosa*, which in turn drives these wounds toward persistent non-resolving inflammation and contributes to impaired healing (*10, 23, 24*). Here, we demonstrate that early after injury, the diabetic wound exhibits a paradoxical and damaging decrease in essential neutrophil trafficking, which in turn renders diabetic wounds vulnerable to infection. Our data point to impaired signaling through the FPR1 chemokine receptor (resulting from exposure to high glucose levels), as an important culprit responsible for the delay in the neutrophil influx in response in diabetic wounds early after injury.

It is worth noting that 1h exposure to high glucose levels dramatically impaired chemotaxis signaling through the FPR1, suggesting that even a short-term rise in serum glucose levels could potentially make non-diabetic people transiently immunocompromised and susceptible to infection. In line with this notion, hyperglycemia during the perioperative and postoperative periods are found to be significant risk factors for surgical site infections (SSIs) (*65, 66*), while glycemic control during the perioperative period has been shown to significantly reduce SSI rates both in human and animals (*44, 66*). It remains unclear why exposure to high glucose dampens the expression and signaling through the FPR1 chemokine receptor. We posit that it may involve metabolic changes, resulting from high glucos in neutrophils. We are actively investigating this possibility.

Our data demonstrate that at least the expression and signaling through CCR1 and CXCR2 auxiliary receptors are not adversely affected by high glucose, but they may not be signaling in diabetic wounds early after injury because of deficiency in the production of their ligands. It remains a possibility that other auxiliary chemokine receptors which amplify the neutrophil migration in wounds and toward infection (e.g, CXCR1, BLT1, etc. (*31*)), may also remain functional under diabetic condition and their engagement with their respective ligands could similarly enhance antimicrobial defenses in diabetic wounds. Future studies should assess these possibilities and evaluate how serum glucose level affects the expression and activity of all the ∼30 chemokine receptors that mediate chemotaxis in neutrophils in diabetic individuals and toward infection.

It is encouraging that one-time topical treatment with CCL3 substantially boosted antimicrobial defenses and stimulated healing in diabetic wounds. However, given that diabetic foot ulcers are already suffering from neutrophilia and heightened inflammation, the therapeutic value of CCL3 treatment may seem questionable. We posit that CCL3 topical treatment may have real therapeutic potential in diabetic wound care, at least in a subset of diabetic individuals represented by our animal model, if applied topically after the surgical wound debridement process. The purpose of surgical debridement, which is performed as a standard-of-care weekly or biweekly in the clinics, is to convert a chronic non-healing wound environment into an acute healing environment through the removal of necrotic and infected tissue, and the senescent and non-responsive cells (*67-69*). Therefore, debrided wound environment is likely to be more similar to day 1 fresh wounds than day 10 chronic wounds in our studies. Future studies are needed to evaluate the therapeutic potential of CCL3 in diabetic wound care.

## Materials and Methods

### Procedures related to animal studies

We have an approval from the Rush University Medical Center Institutional Animal Care and Use Committee (IACUC) to conduct research as indicated. All procedures complied strictly with the standards for care and use of animal subjects as stated in the Guide for the Care and Use of Laboratory Animals (Institute of Laboratory Animal Resources, National Academy of Sciences, Bethesda, MD, USA). We obtained 8-week-old C57BL/6 (normal) and their diabetic littermates, C57BLKS-m *lepr*^*db*^ (db/db) mice from the Jackson Laboratories (Bar Harbor, ME). These mice were allowed to acclimate to the environment for 1 week prior to experimentation. Wounding and wound infection were carried out as we described previously (*10, 25*). Hematoxylin & Eosin (H&E) staining were performed as we described previously (*10, 27*). Neutrophil trafficking into wounds was assessed by immunohistochemical (IHC) analysis using Ly6G staining as described previously (*70*). Wound tissues’ contents of myeloperoxidase (MPO) were assessed by ELISA as described (*27*). CCL3 expression was assessed by RT-PCR, following the protocol we described previously (*25*). To account for reduced neutrophil migration into day 1 diabetic wounds, data were normalized by 18S RNA levels. We used *Pseudomonas aeruginosa* PA103 in these studies. This strain has been described previously (*71, 72*) and we have shown that it causes massive infection and exacerbates wound damage in db/db wounds (*10*). Infection levels in wounds were evaluated by determining the number of bacteria (colony forming unit (CFU)) per gram of wound tissues, as we described (*10, 44*).

### Histological analyses and wound healing assessment

Wound healing was assessed by digital photography; by re-epithelization assessment using H&E staining; by fibroblasts and myofibroblasts tissue content analyses using vimentin and α-SMA; and by elastin and collagen matrix deposition assessment using elastin or Masson’s Trichrome staining, using previously described techniques*(10, 25, 59, 73, 74)*. The histological data, (obtained from n≥5 mice/group and ≥9 random fields/wound/mouse), were normalized per wound surface area.

### Neutrophil isolation from human and mouse

We have an Institutional Review Board (IRB)-approved protocol in accordance with the Common Rule (45CFR46, December 13, 2001) and any other governing regulations or subparts. This IRB-approved protocol allows us to collect blood samples from non-diabetic volunteers with their consents for these studies. The blood samples were first checked by a glucometer kit (FreeStyle Lite, Blood Glucose Monitoring System) to ensure that blood glucose level is within the normal range, ≤ 100 mg/dl. Next, human neutrophils were purified from blood using the EasySep™ Human Neutrophil Enrichment Kit (STEMCELL Technologies), according to manufacturer’s protocol.

Murine neutrophils were isolated from either peripheral blood (used in Fig. 2a-c; Fig. S3a; Fig. 3h-i) or bone marrow (Fig. 2f-m, Fig. 3b-g, and Fig. S3d) for the studies involving glucose exposure using EasySep™ Mouse Neutrophil Enrichment Kit (STEMCELL Technologies), as per manufacture’s protocol and as described previously (*25, 75*). Mouse neutrophils involving comparisons between C57B normal and db/db diabetic neutrophils were extracted from N=4 blood pools/group, with each blood pool being from 4 mice: totaling 16 mice per group. This was to obtain enough neutrophils from mouse blood (∼0.8 ml of blood/mouse, 3.2 ml total) for analyses to achieve statistical significance.

### Neutrophil chemotactic response

Purified human and murine neutrophils were incubated in (IX HBSS with 2% HSA) containing glucose at indicated concentrations for 1h at 37°C and stained with Calcein AM (5ug/mL) for 30 minutes. After washing the cells, the cell migration assay was performed *in vitro* using 96-well disposable chemotaxis chambers (Cat. No. 106-8, Neuro Probe, Gaithersburg, MD, USA). Neutrophils chemotaxis toward the chemoattractants (chemokines) were performed at indicated concentrations, or PBS (to account for the background neutrophil migration), following the manufacturer’s protocol. Cell migration was assessed by a fluorescence (excitation at 485nm, emission at 530nm) plate reader Cytation 3 Cell Imaging Multi-Mode Reader (Biotek Instruments, Inc). The actual chemotaxis values were obtained by subtracting random chemotaxis values (PBS) from the chemotaxis values in response to chemokines.

### Flow cytometry

#### Wound tissue digestion and flow cytometric

C57B and db/db wound tissues were obtained at indicated timepoints as described (*25*), weighed, and place immediately in cold HBSS (Mediatech, Inc., Manassas, VA). Subcutaneous fat was removed using a scalpel and scissors were used to cut the tissue into small < 2mm pieces. The tissue was enzymatically dissociated in DNAse I (40µg/ml; Sigma-Aldrich Co., St. Louis, MO) and Collagenase D (1mg/ml HBSS; Roch Diagnostics, Indianapolis, IN) at 37°C for 30 minutes. Cold PBS was used to stop the dissociation process. The tissue was then mechanically dissociated using the gentleMACS octoDissociator (program B; Miletynyi Biotec, Auburn, CA) and passed through 70µm nylon screens into 50ml conical tubes. Cells were washed twice with PBS. Resultant single-cell suspensions were stained using the indicated fluorescently labeled antibodies against cell surface markers, according to standard protocols described previously (*76, 77*). All antibodies were purchased from eBioscience, Inc. (San Diego, CA). Flow cytometry was performed using a the LSRFortessa cell analyzer (Becton, Dickinson, and Company)) and data were analyzed using FlowJo software (Tree Star, Ashland, OR), as previously described (*25, 78*). Briefly, for the gating strategy, Live singlet lymphocytes were identified by gating on forward scatter-area (FSC-A) versus (vs.) side scatter-area (SSC-A), then LIVE/DEAD staining vs. SSC-A, FSC-A vs. FSC-height (H), SSC-A vs. SSC-H, FSC-width (W) vs SSC-W, and CD45 vs SSC-A. T cells, B cells, and NK cells were excluded using antibodies against CD3, CD19, and NK1.1, respectively, all on one channel as a dump gate. Neutrophils were then identified using CD11b vs Ly6G staining, with neutrophils being CD11b high and Ly6G high. FPR1 and CCR1 expression on neutrophils was then analyzed and is presented as percentage of cells (e.g., neutrophils) expressing the respective marker.

#### Neutrophil depletion in mice

Neutrophil depletion in mice were performed as described (*58, 79*). Briefly, db/db mice received either anti-Ly6G (100µg/mouse) to cause neutrophil depletion or an IgG isoform control (100µg/mouse), by intraperitoneal (i.p.) injection. Neutrophil depletion was confirmed by the assessment of neutrophil content in the blood (circulation) by flow cytometry or in wound tissues by MPO analysis.

#### Western blot analyses

We performed Western immunoblotting on cell lysates or on tissue lysates, using the indicated antibodies as we described previously (*27, 71, 80*). Equal amounts of proteins (as determined by BCA analysis) were loaded. GAPDH was used as a loading control.

#### Gene expression analysis by Real-Time Polymerase Chain Reaction (RT-PCR)

Gene expression was assessed by RT-PCR as we described (*25*): cDNA was generated using SuperScript™ III First-Strand Synthesis System cDNA Synthesis Kit (Cat. No. 18080051) from Thermo Fisher, according to manufacturer’s protocol. RT-PCR was then preformed with gene-specific primer pairs mentioned below, using the Applied Biosystems QuantStudio™ 7 Flex Real-Time PCR System. The data were calculated using the 2^−ΔΔCt^ method and were presented as ratio of transcripts for gene of interest normalized to 18S. We performed RT-PCR using the following primers: FPR1 forward: GAGCCTAGCCAAGAAGGTAATC, reverse: TCCCTGGTCCAAGTCTACTATT; Phospholipase C gamma 1 (Plcg1) forward: GGTGAGGCCAAATGTGAGATA, reverse: GGGCAACCAAGAGGAATGA; Chemokine (C-C motif) receptor 1 (Ccr1) forward: GCTATGCAGGGATCATCAGAAT, reverse: GGTCCAGAGGAGGAAGAATAGA; Chemokine (C-C motif) ligand 3 (Ccl3) forward: TCACTGACCTGGAACTGAATG, reverse: CAGCTTATAGGAGATGGAGCTATG; 18S forward: CACGGACAGGATTGACAGATT, reverse: GCCAGAGTCTCGTTCGTTATC.

#### Antibodies (for neutrophil depletion study)

Anti-Ly6G monoclonal antibody clone RB6-8C5 (Cat. No MA1-10401 from Invitrogen, Mouse (G3A1) mAb IgG1 Isotype Control #5415 (Cell Signaling Technologies).

#### Antibodies (for IHC and Western blotting)

Anti-Ly6G antibody clone RB6-8C5 for IHC (#ab25377 from Abcam); anti-FPR1 (Cat. No. NB100-56473 from NOVUS Biological); anti-PLCϒ1 (Cat. No. cs2822 from Cell Signaling), GAPDH Antibody Rabbit Polyclonal, Cat. No. 10494-1-AP (Proteintech Cat. No. 1094-I-AP); and CCR1 Polyclonal antibody (Abnova Cat. No. PAB0176). Anti-α-SMA antibody (Cat. No. ab5694) and anti-vimentin antibody (Cat. No. ab92547) from Abcam.

#### Antibodies (for flow cytometry)

Mouse CCR1 Alexa Fluor® 488-conjugated Antibody #FAB5986G-100UG (R & D Systems); Alexa Fluor® 700 anti-mouse NK-1.1 Antibody #108729 (BioLegend); Alexa Fluor® 700 anti-mouse CD3ε Antibody #152315 (BioLegend); Alexa Fluor® 700 anti-mouse CD19 Antibody #115527 (BioLegend); BV605 Hamster Anti-Mouse CD11c Clone HL3 (RUO) #563057 (BD Biosciences); LIVE/DEAD™ Fixable Aqua Dead Cell Stain Kit, for 405 nm excitation #L34966 (ThermoFisher Scientific); F4/80 antibody | Cl:A3-1 #MCA497PBT (Bio-Rad); BV650 Hamster Anti-Mouse CD11c Clone HL3 (RUO) #564079 (BD Biosciences); BV711 Rat Anti-Mouse CD45 Clone 30-F11 (RUO) #563709 (BD Biosciences); NK1.1 Monoclonal Antibody (PK136), PE, eBioscience™ #12-5941-82 (ThermoFisher Scientific); CD19 Monoclonal Antibody (eBio1D3 (1D3)), PE, eBioscience™ #12-0193-82 (ThermoFisher Scientific); CD3e Monoclonal Antibody (145-2C11), PE, eBioscience™ #12-0031-82 (ThermoFisher Scientific); FPR1 Polyclonal Antibody #PA1-41398 (ThermoFisher Scientific); Goat anti-Rabbit IgG (H+L) Highly Cross-Adsorbed Secondary Antibody, Alexa Fluor 594 #A-11037 (ThermoFisher Scientific); Ly6G Monoclonal Antibody (1A8-Ly6g), PE-Cyanine7, eBioscience™ #25-9668-82 (ThermoFisher Scientific), PerCP Cy5.5 CD45 (BD Biosciences, Cat. 550994); APC Gr1, PE CD11b (BD Biosciences, Cat 553129); FITC CD69 (BD Biosciences, Cat #: 557392); and PECy7 F4/80 (Biolegends, Cat 123114).

#### Reagents

Hematoxylin & Eosin Staining (Richard Allan Scientific Hematoxylin, Eosin Y, and Bluing Reagent Cat. Numbers: 7111L, 7211L, and 7301L from Thermo Fisher; Myeloperoxidase (MPO) Mouse ELISA Kit; cAMP measurement by ELISA (Cyclic AMP Competitive ELISA Kit); CCL3 (recombinant mouse CCL3/MIP-1α protein; 450-MA, rhCCL3/MIP-1α isoform LD78a; 270-LD from R&D); N-formyl-Met-Leu-Phe (fMLF), Cat. No. 59880-97-6 from Sigma; Collagenase D (CAS No. 9001-12-1 from Sigma Aldrich); Masson’s Trichrome (Trichrome Stain Connective Tissue Stain; Cat. No. ab150686 from Abcam). EasySep Human neutrophil Enrichment Kit, EasySep Mouse neutrophil Enrichment Kit, EasySep Buffer (Cat. No. 19762, 19666, and 20144 from STEMCELL Technologies), and Calcein AM (Cat no. C1430 from ThermoFischer).

#### Statistical analysis

Statistical analyses were performed using GraphPad Prism 6.0 as we described previously (*81, 82*). Comparisons between two groups were performed using Student’s *t*-test. Comparisons between more than two groups were performed using one-way analysis of variance (one-way ANOVA). To account for error inflation due to multiple testing, the Bonferroni method was used. Data are presented as Mean ± SEM. Statistical significance threshold was set at *P*-values ≤ 0.05.

## Acknowledgments

We are thankful to Dr. Lena Al-Harthi and Dr. Celeste Napier for the use their equipment. We also would like to thank Mr. Jeffrey Martinson for his help with the flow cytometry and the rest of Shafikhani lab for their valued opinions on these studies. This work was supported by the National Institutes of Health (NIH) grant RO1DK107713 to (S.H.S.), R01AI150668 to (S.H.S.), F31DK118797 to (J.Z.), and the NIH PhD institutional training grant GM109421.

## Author contributions

S. H. S. conceived and coordinated all aspects of the studies and wrote the paper. R.R & J.Z. conducted and contributed to all the figures in this MS and wrote the paper. S.J.W. contributed to Figure 2. M.F.M. contributed to animal studies. A.Z., R.E., & E.F.G. contributed to the flow cytometry data in all figures. T.K. and J.R. contributed to the data analyses, research design, and reagents. All authors had the opportunity to review and edit the manuscript. All authors approved the final version of the manuscript.

---

**Fig. S1.**
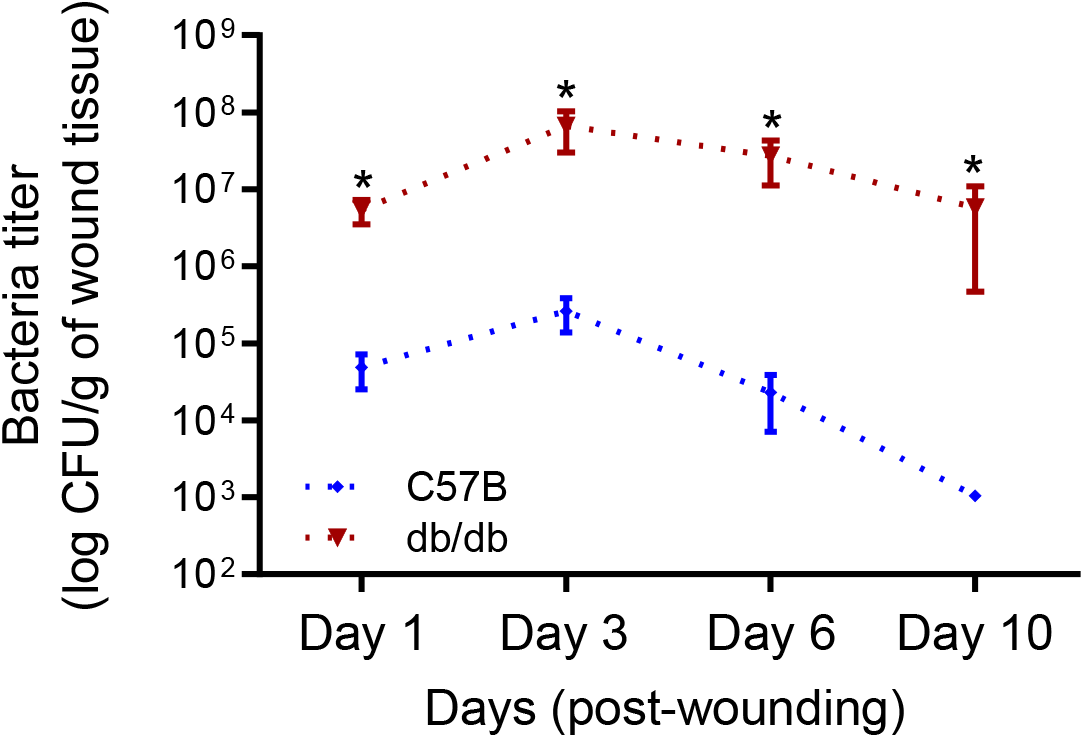
Diabetic wound is vulnerable to increased infection with *Pseudomonas aeruginosa*. Normal and diabetic wounds were infected with 10^3^ of *P. aeruginosa* (PA103). Bacterial burden in wounds was determined by serial dilution and plating at indicated times after infection and is shown as the Mean ± SEM. (N=8; 4 mice/group, 2 wounds/mouse; (*) Represents significance with p<0.01. Statistical analyses between groups were conducted by One-way ANOVA with additional post hoc testing, and pair-wise comparisons between groups were performed or by unpaired Student’s *t*-test).

**Fig. S2.**
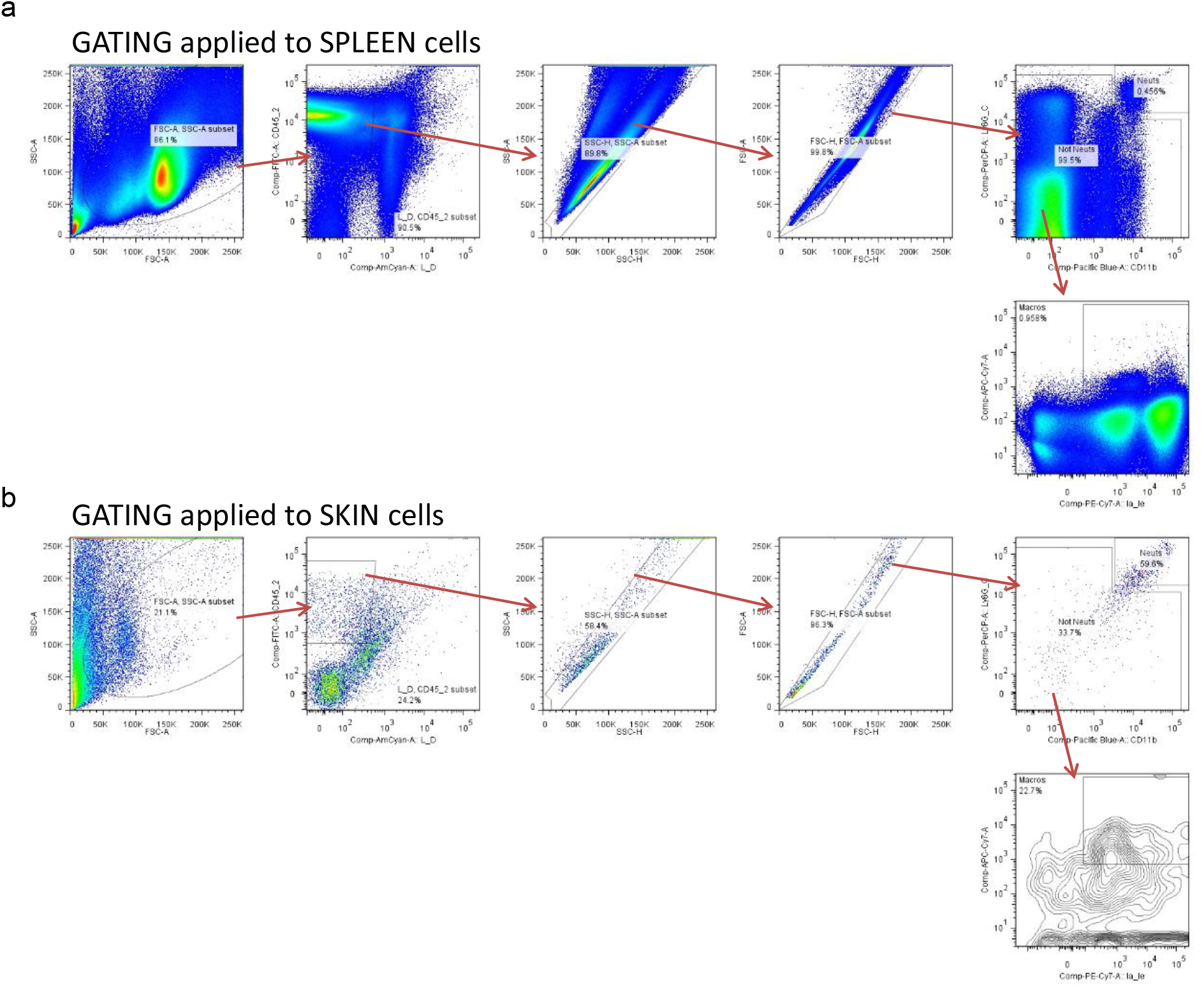
Gating strategy for flow cytometric analysis. Spleen **(a)** and skin tissues **(b)** were harvested from C57B mice. For the gating strategy, Live singlet lymphocytes were identified by gating on forward scatter (FSC)-area (A) versus (vs) side scatter (SSC)-A, then LIVE/DEAD staining vs SSC-A, FSC-A vs FSC-height (H), SSC-A vs SSC-H, FSC-width (W) vs SSC-W, and CD45 vs SSC-A. T cells, B cells, and NK cells were excluded using antibodies against CD3, CD19, and NK1.1, respectively, all on one channel as a dump gate. Neutrophils were then identified using CD11b vs Ly6G staining, with neutrophils being CD11b high and Ly6G high. Macrophages were identified as CD11b positive and Ly6G low/negative, followed by F4/80 positive staining.

**Fig. S3.**
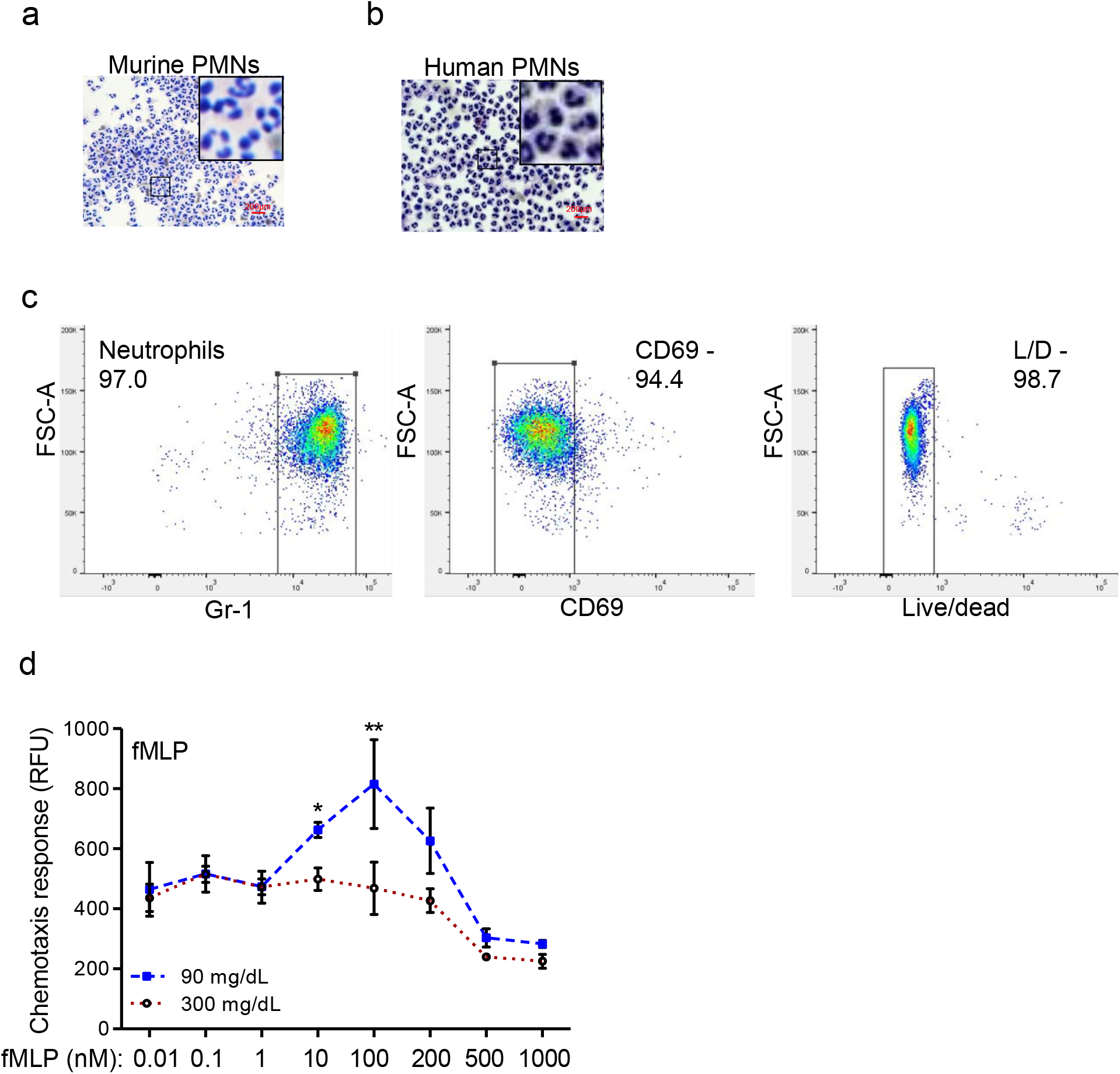
Chemotactic response is impaired in diabetic neutrophils through FPR1 primary receptor. (**a-b**) Neutrophils (PMNs) were purified from murine (C57B bone marrow) and human peripheral blood, as discussed in Materials and Methods. Representative images of mouse and human purified neutrophils are shown at indicated magnification. Magnified representative regions are shown inserts within each image. (Red scale bars are 200μm). (**c**) Representative flow histograms of purified mouse neutrophils showing that these neutrophils are over 97% pure, live, and naïve, as assessed by indicated markers. (**d**) Chemotaxis of purified mouse PMNs towards varying concentrations of fMLP after 1h exposure to normal glucose (90 mg/dl) or high glucose in diabetic range (300 mg/dl). Data are plotted as the Mean ± SEM. (N=4 mice/group; ns = not significant; *p<0.05, **p<0.01, ***p<0.001. Statistical analyses between groups were conducted by One-way ANOVA with additional post hoc testing, and pair-wise comparisons between groups were performed or by unpaired Student’s *t*-test).

**Fig. S4.**
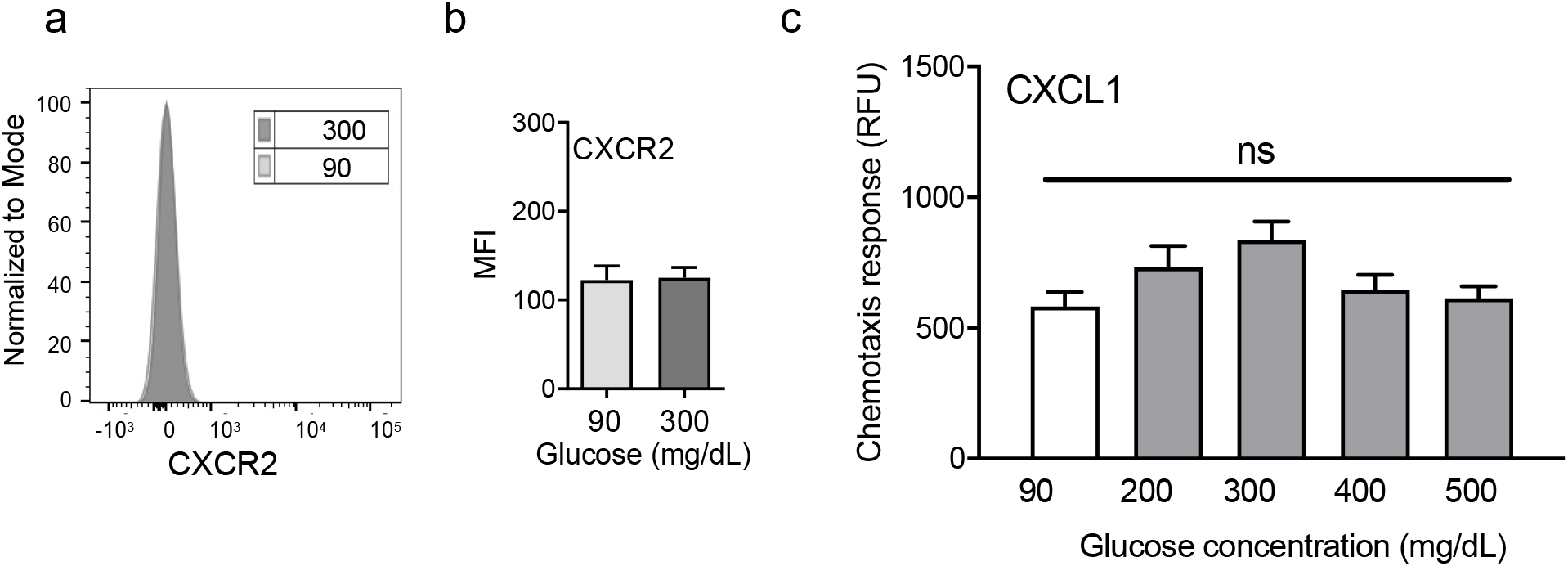
Exposure to high glucose does not affect CXCR2 auxiliary chemokine receptor. **(a-b)** Mouse neutrophils were exposed to glucose at indicated concentrations for 1h and evaluated for their surface expression of CXCR2 by flow cytometry. A representative histogram is shown in **(a)** and the corresponding data are plotted as the Mean ± SEM is shown in **(b)**. (N=4). **(c)** Murine neutrophils were examined for their chemotactic response toward CXCL1 (5ng/ml) and after 1h exposure to normal glucose (90 mg/dl) and high glucose in diabetic range (200-500 mg/dl). Data were plotted as Mean ± SEM. (N=6; ns = not significant).

**Fig. S5.**
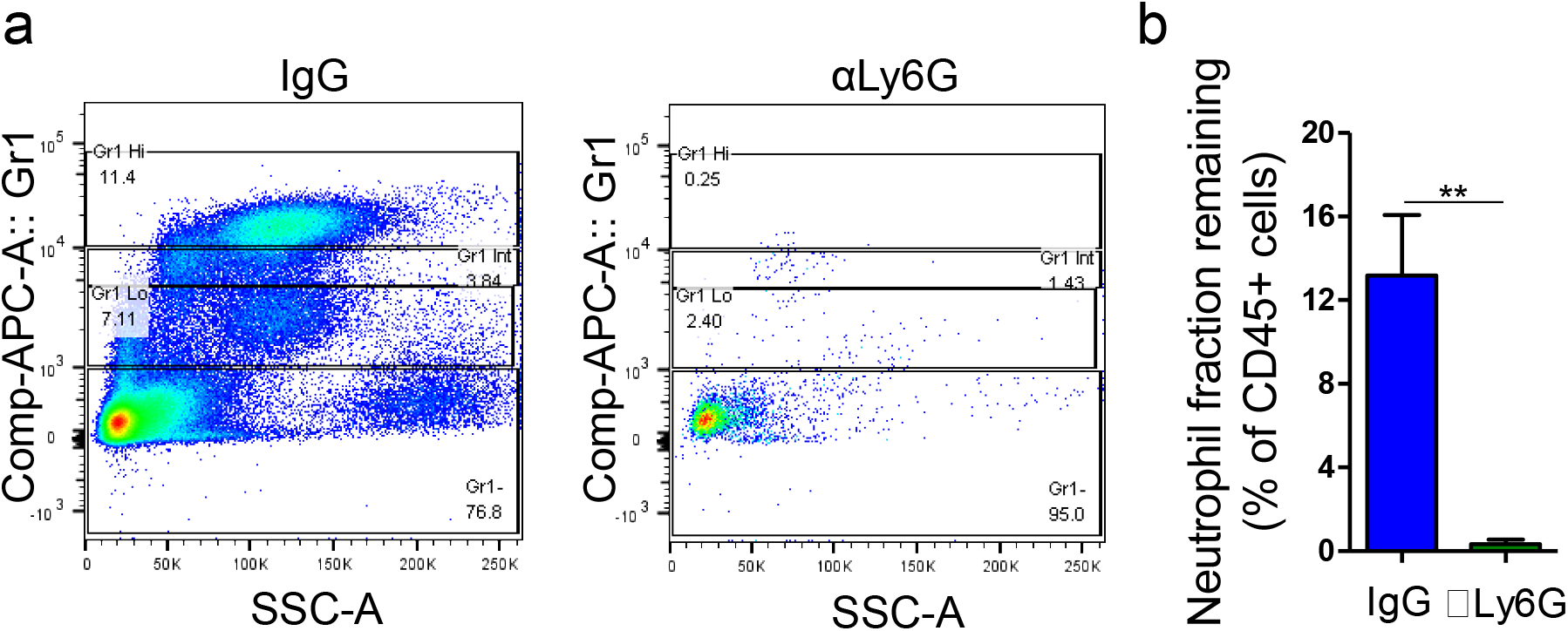
Supplementary data associated with Fig. 4. db/db mice were injected by i.p with anti-Ly6G or IgG isoform. 24h after injection, their peripheral bloods were examined for their neutrophil contents by flowcytometry. Representative histograms of neutrophil depletion are shown in (**a**) and the corresponding data plotted as the Mean ± SEM is shown in (**b**). (N=4; **p<0.01. Student’s *t*-test).

**Fig. S6.**
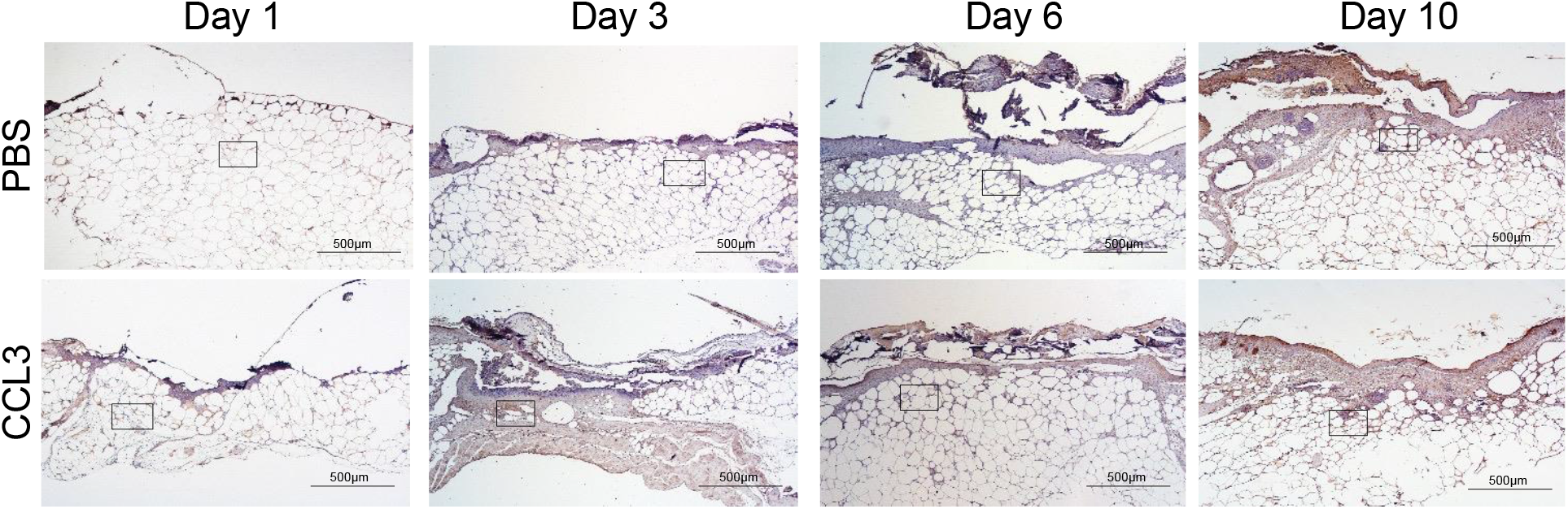
Full wound images associated with Fig. 5c. db/db animals were wounded and treated with either CCL3 or PBS prior to infection with PA103 (10^3^ CFU). 24h after treatment and infection, wound tissues were harvested and stained with neutrophil marker Ly6G. Representative low magnification (40X) images of full wounds are shown. Inserted rectangles show the cropped regions represented in Fig 5c. (Black scale bar = 500µm).

**Fig. S7.**
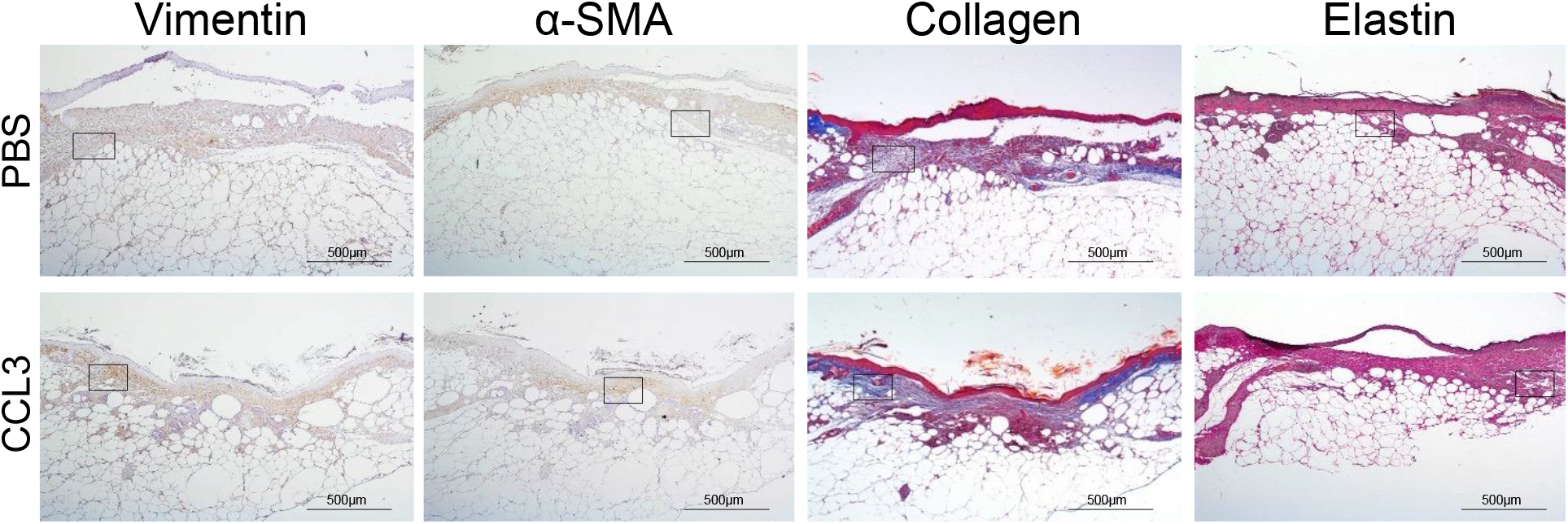
Full wound images associated with Fig. 6e. db/db animals were wounded and treated with either CCL3 or PBS prior to infection with PA103 (10^3^ CFU). 10 days after treatment and infection (Day 10), wound tissues were harvested and assessed for fibroblast, myofibroblast, elastin and cartilage healing markers by vimentin, α-SMA, Masson’s Trichrome and elastin staining respectively. Representative 40X magnification images of the full wounds are shown, and the high magnification images and the tabulated data are presented in Fig. 6e-f. (Black scale bar = 500µm. Inserted rectangles show the cropped regions represented in Fig. 6e).

